# Conformational flexibility of soybean lipoxygenase is coupled to crystal solvent content in serial crystallography

**DOI:** 10.64898/2026.05.30.729005

**Authors:** Alexander M. Wolff, Daniel W. Paley, Iris D. Young, Alexander Deary, Vidya Ganapati, Jack Hirschman, Azeem Horani, Randy Lemons, Stella Lisova, Ralph L. McAnelly, Dondis Moreland, Sandra T.M. Mous, Amanda Ohler, Joshua M. Rodriguez, Silvia Russi, Raymond G. Sierra, Daniel Mariusz Tchoń, Jan Paulo T. Zaragoza, Sergio Carbajo, Alec H. Follmer, Judith P. Klinman, Adam R. Offenbacher, Frédéric Poitevin, Nicholas K. Sauter, Mark A. Wilson, Aaron S. Brewster, Michael C. Thompson

**Affiliations:** Department of Chemistry and Biochemistry, University of California, Merced, Merced, CA, USA; Molecular Biophysics and Integrated Bioimaging Division, Lawrence Berkeley National Laboratory, Berkeley, CA, USA; Linac Coherent Light Source, SLAC National Accelerator Laboratory, Menlo Park, CA, USA; Department of Chemistry, East Carolina University, Greenville, NC, USA; Stanford Synchrotron Radiation Lightsource, SLAC National Accelerator Laboratory, Menlo Park, CA, USA; Department of Chemistry, University of California Berkeley, Berkeley, CA, USA; Department of Electrical and Computer Engineering, University of California, Los Angeles, Los Angeles, CA, USA; Department of Chemistry, University of California, Davis, Davis, CA, USA; Department of Biochemistry and Redox Biology Center, University of Nebraska, Lincoln, NE, USA

**Keywords:** serial crystallography, protein dynamics, indexing ambiguity, polymorphism

## Abstract

X-ray crystallography is increasingly employed to study protein conformational ensembles under physiological conditions, but the effects of the crystal lattice on protein motions remains understudied. Here, we report the structure determination of soybean lipoxygenase-1 (SLO) from microcrystal slurries using serial femtosecond crystallography (SFX) at the Linac Coherent Light Source. During data analysis, we observed unexpected polymorphism in SLO’s unit-cell parameters, arising from two compounding factors: indexing ambiguities caused by the pseudo-tetragonal symmetry of the SLO crystal lattice, and true non-isomorphism between individual crystal populations consistent with different solvent content. By combining unit-cell clustering with systematic reindexing, we resolved two distinct polymorphs and determined two independent structures from a single experiment. The two structures exhibit a small overall RMSD (0.34 Å), yet a difference distance matrix reveals coordinated rearrangements that are not readily apparent from simple structural overlays. Furthermore, a difference of approximately 8.5% in crystal solvent content produces measurable differences in crystal contacts and conformational flexibility. The more hydrated (large-cell) polymorph exhibits greater inter-domain flexibility, as well as higher B-factors in key hydrophobic core residues. In the dehydrated (small-cell) polymorph, which shows less interdomain flexibility, these same residues adopt discrete alternative conformations resolvable in the electron density. Our results highlight that subtle changes in crystal packing can give rise to distinct conformational landscapes for crystallized proteins, with potential implications for the interpretation of protein intramolecular dynamics from crystallographic data.

**Synopsis:** Using soybean lipoxygenase-1 as a model system, we determine two independent structures within a single experiment that differ in crystal solvent content. Despite a small overall RMSD, a detailed analysis reveals coordinated domain-level rearrangements and shows that conformational flexibility is sensitive to crystal packing, a consideration of broad relevance for the study of protein dynamics using X-ray crystallography.

## 1. Introduction

The role of solvent content in determining the behavior of protein crystals has been known for essentially as long as the field of macromolecular crystallography has existed. In order to collect the first ever diffraction images from protein crystals, Bernal and Crowfoot made the key discovery that their crystals only produced diffraction patterns if they remained hydrated (Bernal & Crowfoot, 1934). Later, Crick and Kendrew noted that most protein crystals have a solvent content of 40-60% (Crick & Kendrew, 1957), an observation that was roughly in agreement with a later study by Matthews relating solvent content to the packing density of protein in the asymmetric unit (Matthews, 1968). As the volume of data in the Protein Data Bank grew, Rupp and colleagues showed that there was a correlation between the solvent content of crystals and the resolution of the resulting diffraction patterns, with lower solvent content generally yielding higher resolution (Kantardjieff & Rupp, 2003; Weichenberger *et al*., 2015). This observation spawned numerous attempts to improve the quality of X-ray diffraction from crystals by modulating their solvent content (Bowler *et al*., 2006; Sanchez-Weatherby *et al*., 2009; Bowler *et al*., 2015; Russi *et al*., 2011; Heras & Martin, 2005). In a narrower range of contexts, the relationship between crystal solvent content and protein conformation has been explored. Notably, changes in solvent content have been associated with changes to backbone and side chain conformation, as well as changes to hydration layer structure, in lysozyme (Kodandapani *et al*., 1990; Madhusudan *et al*., 1993; Nagendra *et al*., 1998; Biswal *et al*., 2000; Atakisi *et al*., 2018) and bovine pancreatic ribonuclease A (Bell, 1999). Indeed, it is well established that changes in solvent content can convert a protein crystal between different polymorphs, affecting the quality of X-ray diffraction data and the structures of the crystallized proteins. On the other hand, the effect of solvent content and crystal packing on protein motions within the crystal lattice remains understudied.

Questions about the effect of the crystal environment on protein dynamics have come into focus for serial crystallography experiments over the past decade, particularly as the method is increasingly applied to study functionally important protein motions using time-resolved approaches. In a serial crystallography experiment, thousands of diffraction images comprising a dataset are collected from distinct crystal specimens, typically using batch-grown microcrystals that are no larger than tens of microns in each dimension (Chapman *et al*., 2011; Boutet *et al*., 2012; Schlichting, 2015). These microcrystal samples are frequently delivered to the X-ray beam using microfluidic jet-based systems that expose them to high pressure (Weierstall *et al*., 2014), high voltage (Sierra *et al*., 2012), or vacuum (DePonte *et al*., 2008), conditions which can physically damage and disrupt microcrystals (Martiel *et al*., 2019). Additionally, samples are often mixed with solutes that act as viscogens, including water-soluble polymers such as polyethylene glycol and cellulose derivatives (Kovácsová *et al*., 2017; Sugahara *et al*., 2017), to alter their rheological properties and stabilize their flow. Experimentalists often must test several conditions before settling on the most appropriate one, and choices can be limited by the properties of the sample and the requirements of the experiment. Sample parameters (e.g. viscogen concentration) and microfluidic parameters (e.g. pressure), are also typically manipulated “on-the-fly” to empirically optimize data collection. Considerations regarding sample delivery further complicate questions about the relationship between solvent content, structure, and dynamics of crystallized molecules, because treatments and manipulations that are often required for sample delivery are known to modify the solvent content of crystals and the behavior of the crystallized molecules in the context of single crystal experiments (Collins *et al*., 2011; Atakisi *et al*., 2018; Heras & Martin, 2005; Hekstra *et al*., 2016). Perturbations associated with sample delivery in serial crystallography experiments have not been systematically studied, but their effects are likely not negligible.

Here we report the results of a SFX experiment on SLO, which illustrate complications that can arise from crystal polymorphism coupled to differences in solvent content, and highlight how these differences can alter the flexibility of the crystallized molecules. SLO is an ideal system to study using SFX. It is a non-heme iron metalloenzyme (Roza & Francke, 1973; Chan, 1973; Dunham *et al*., 1990), so the ability of ultrafast X-ray pulses to outrun the effects of radiation damage preserves the chemistry of the active site during structural measurements (Chapman *et al*., 2014; Weierstall *et al*., 2014). Furthermore, solvent-coupled protein dynamics have been associated with the catalytic mechanism of SLO (Zaragoza *et al*., 2023; Offenbacher *et al*., 2017a; Zaragoza *et al*., 2019), making it an interesting candidate for time-resolved experiments, which are facilitated by the serial format of data collection (Follmer *et al*., 2025). When performing SFX experiments on SLO, we encountered unexpected polymorphism in SLO’s unit-cell parameters. Analysis revealed the polymorphism was caused by a combination of indexing ambiguities related to the pseudosymmetry of the SLO crystal lattice, and true non-isomorphism between individual crystals, consistent with differences in crystal solvent content. Sorting these crystal polymorphs allowed us to determine two unique structures of SLO from a single experiment. The refined structures align with a global RMSD of 0.34 Å, but this apparent similarity belies differences in the enzyme’s conformational landscape that only became evident after a careful analysis of anisotropic B-factors (refined using a TLS model) and alternate side chain conformations. These differences were notable for hydrophobic residues in the protein core which have previously been shown to be important for SLO’s catalytic activity (Offenbacher *et al*., 2017b). Our work on SLO illustrates a general challenge for the interpretation of protein motions from crystallographic data: unit cell distortions can introduce differences in protein dynamics across crystal polymorphs.

## 2. Materials and Methods

### 2.1 Expression and Purification of SLO

SLO was expressed and purified as previously described (Hu *et al*., 2014; Offenbacher *et al*., 2017b). Briefly, the plasmid containing an untagged, wild-type SLO gene was transformed into BL21(DE3) Codon Plus RIL cells. An overnight LB starter culture, supplemented with ampicillin and 0.4% glucose, was grown overnight with shaking at 30 °C (Jakobowski *et al*., 2024). The overnight was added to 2×YT media (100-fold) containing AMP and 3% ethanol, and grown at 37 °C to an OD600 of 1.0-1.2. The temperature of the shaker was reduced to 18 °C and the cells were collected after 48 hours. The cells were resuspended in a lysis buffer and sonicated. The SLO protein was exchanged in 20 mM Bis-Tris, pH 6.0 and isolated in two steps with two cation exchange columns (SP Sepharose and UNO S6 [Bio-Rad]). For each column, the sample was loaded and washed with 2 column volumes of 20 mM Bis-Tris,pH 6, followed by elution with a linear gradient (0-500 mM NaCl in 20 mM Bis-Tris buffer, pH 6). The eluted protein was dialyzed in 0.1 M borate, pH 9 buffer and stored at -80 °C until further use. The protein, as isolated, was estimated to contain approximately 0.8 iron atoms per protein in the ferrous state.

For crystallography, additional purification steps were performed. SLO (from above) was thawed and dialyzed for three hours in 50 mM sodium acetate (pH 5). Aggregation was removed by centrifugation and the protein was further purified with a HiPrepTM 26/60 SephacrylTM S-200 HR column equilibrated in 50 mM HEPES, 150 mM NaCl (pH 7.4). Protein samples were stored at -80°C until further use. Upon thawing, protein was immediately dialyzed back into 50mM sodium acetate (pH 5) prior to crystallization.

### 2.2 Seed Stock Preparation

SLO at 5 mg/mL in 50 mM NaOAc, pH=5.0, was mixed in a 1:1 ratio with mother liquor (0.4M NaOAc pH=5.5, 8-10% PEG-3350) in 2 µL hanging drops. Crystals formed within 5-7 days (Steczko *et al*., 1990). Crystals from a single hanging drop were crushed, then diluted to ∼50 uL with mother liquor. This seed stock was used to make the first batch of microcrystals. Subsequent seed stocks were prepared by taking ∼10µL from previous batch crystal solutions, crushing the crystals, then diluting with mother liquor to ∼50 µL.

### 2.3 Batch Crystallization

SLO at 10 mg/mL in 50 mM NaOAc, pH=5.0, was mixed in a 1:1 ratio with batch precipitant (0.4M NaOAc pH=5.5, 16% PEG-3350) to a volume of 1 mL in a 1.5 mL Eppendorf Tube. ∼10µL of seed stock (see 2.2.1.) was added to the tube cap. The tube was then placed on a Thermo Fisher Tube Revolver Rotator, set to 30 RPM. Samples were left to mix overnight. When ready for use, crystals were allowed to settle at the bottom of the Eppendorf Tube, and excess crystallization buffer was removed. Crystals were mixed in a ∼1:1 ratio with 18% (w/v) hydroxyethylcellulose (Sigma-Aldrich PN-09368) dissolved in crystallization buffer (0.225M NaOAc pH=∼5.43, 8% PEG-3350). Each component was loaded into a Hamilton Syringe, and both syringes, along with a third empty syringe, were connected to a custom 3-way coupling device (James *et al*., 2019). The batch crystals and cellulose were mixed together and immediately deposited in reservoirs for use with the HVE (high viscosity extrusion) injector (Weierstall *et al*., 2014), with sample injection and data collection typically beginning within 30 minutes of deposition.

### 2.4 Serial Crystallography Data Collection

SFX data were collected at the Macromolecular Femtosecond Crystallography (MFX) endstation of Linac Coherent Lightsource (LCLS) using the Rayonix MX340-HS detector in 2×2 binning mode. Crystals were delivered to the X-ray interaction point using an HVE injector, and data were collected using a ∼5 µm beam at ∼9.5 keV delivered at 30 Hz, with a pulse duration of ∼30 fs. An energy scan was collected across a range of +/- 0.05 keV while monitoring the energy spectra using a crystal transmissive spectrometer (Zhu *et al*., 2013) upline at the XPP instrument (Chollet *et al*., 2015), which allowed for per-shot energy calibration. Detector geometry was calibrated using silver behenate powder diffraction patterns. Further parameters are provided in Table 1.

**Table 1.** Sample delivery and data collection parameters.

|  |  |
| --- | --- |
| Delivery Matrix | 18% hydroxyethyl cellulose |
| Sample flow rate ($\mu\text{L}/\text{min}$) | 0.5 |
| Capillary diameter ($\mu\text{m}$) | 75 |
| Linear jet velocity ($\text{mm}/\text{s}$) | 1.89 |
| Pressure (psi) | ~3200 |
| X-ray Source | LCLS - MFX |
| Photon Energy (keV) | 9.5 |
| X-ray pulse duration (fs) | ~30 |
| Photons per pulse | $\sim 1 \times 10^{12}$ |
| X-ray Repetition Rate (Hz) | 30 |

**Table 2.** Data processing statistics^a^.

|  |  |  |
| --- | --- | --- |
| No. of images | 442472 |  |
| Hits | 65813 → 14.67% |  |
| Indexed | 56902 → 86.46% |  |
| Dataset | Small Cell | Large Cell |
| Lattices Reindexed | 10181 | 34631 |
| Resolution Range | 19.93-1.95 (1.98-1.95) | 19.59-1.95 (1.98-1.95) |
| Space Group | P12<sub>1</sub>1 | P12<sub>1</sub>1 |
| Unit-cell params |  |  |
| a (Å) | 92.04+/-1.12 | 95.97+/-0.31 |
| b (Å) | 93.08+/-0.96 | 94.56+/-0.23 |
| c (Å) | 49.03+/-0.23 | 50.53+/-0.10 |
| α (°) | 90 | 90 |
| β (°) | 92.73 | 91.17 |
| γ (°) | 90 | 90 |
| Total Reflections | 5656883 (11330) | 5099225 (39552) |
| Multiplicity | 94.00 (3.77) | 77.59 (12.19) |
| Completeness (%) | 99.06 (87.53) | 99.88 (99.78) |
| Mean I/sigma (I) | 5.068 (0.622) | 4.416 (0.911) |
| Wilson B (Å<sup>2</sup>) | 23.15 | 32.41 |
| R-split (%) | 12.1 (80.5) | 10.9 (61.8) |
| CC-int (%) | 99.1 (47.1) | 98.9 (56.7) |

<sup>a</sup>Statistics for the highest-resolution shell are shown in parentheses.

### 2.5 Diffraction Data Processing

Data collection was monitored and processed using the *cctbx.xfel* software suite, based on DIALS and CCTBX (Sauter *et al*., 2013; Brewster *et al*., 2025). Spot-finding and initial indexing were carried out using *dials.stills_process*. Images were indexed in space group *P*1 using a target unit cell of (96.03, 94.55, 50.54, 90, 91.15, 90). Initial indexing results were clustered using the *uc_metrics.dbscan* command, which leverages the DBSCAN algorithm (Ester *et al*., 1996) to cluster cells based on values for α, β, and the c-axis. The hyperparameter ε, representing a length scale, was optimized to ensure maximum capture of cells while prioritizing separation of clusters. For ease of separation without losing data, we chose an iterative approach to separate the large cell data. The cells corresponding to Cluster 0 (Fig. 3C) were isolated and the process was repeated on those data alone, yielding four separate clusters (not shown in Fig. 3C) which corresponded to the large unit cell volume (Fig. 3C). Clusters with common c-axis lengths (corresponding to a common unit-cell volume) were considered a group when reindexing (clusters 1,2,3 and 6 from Fig. 3C correspond to the small cell and the four analogous clusters from iterative clustering on cluster 0 from Fig. 3C correspond to the large cell). For a given set of cells assigned to a cluster, reindexing operators were applied to ensure that the mean value of β was >90°. Symmetry constraints appropriate to space group P12_1_1 were then applied, forcing α and ɣ to 90°. Data were then subjected to the Brehm-Diederichs method (Brehm & Diederichs, 2014) to break polar ambiguities using the *modify_cosym* (Gildea & Winter, 2018) module within *cctbx.xfel.merge* (Brewster *et al*., 2025). Finally, these data were merged and postrefined, with error estimates treated using the Ev11 method (Brewster *et al*., 2018; Evans, 2011). Thus, from one stream of data two independent datasets with different unit cell volumes were produced.

### 2.6 Model Building and Refinement

Intensities as output from *cctbx.xfel.merge* were phased via molecular replacement using Phaser (McCoy *et al*., 2007), with chain A from PDB entry 5t5v as a search model. R-free flags were generated and random atomic displacements (0.5 Å) were applied to minimize model bias prior to initial refinement. The model was refined in PHENIX (Afonine *et al*., 2012), beginning with a rigid-body strategy. This was followed by manual rebuilding using Coot (Emsley & Cowtan, 2004) and subsequent refinement in PHENIX. This iteration continued until the model converged. The models and structure factors were deposited to the PDB under codes 9O4S & 9O4T. Statistics for refinement are provided in Table 3. TLS refinement was applied to both models, with 1 TLS group used for the small-cell structure and 3 TLS groups used for the large-cell structure. Contributions to TLS ellipsoids were separated using ECHT (Pearce & Gros, 2021), restricting the model to chain, secondary structure, and atomic levels (auto_levels=chain+ss) with the remaining parameters set to default values. The final structural models were run through the PISA server (Krissinel & Henrick, 2007) to analyze crystal contacts and quantify buried surface area. To systematically evaluate side-chain conformational polymorphism, 2mF_o_-DF_c_ electron density was sampled around the side-chain dihedral angles (e.g., chi_1_, chi_2_) using Ringer (Lang *et al*., 2014).

**Table 3.**
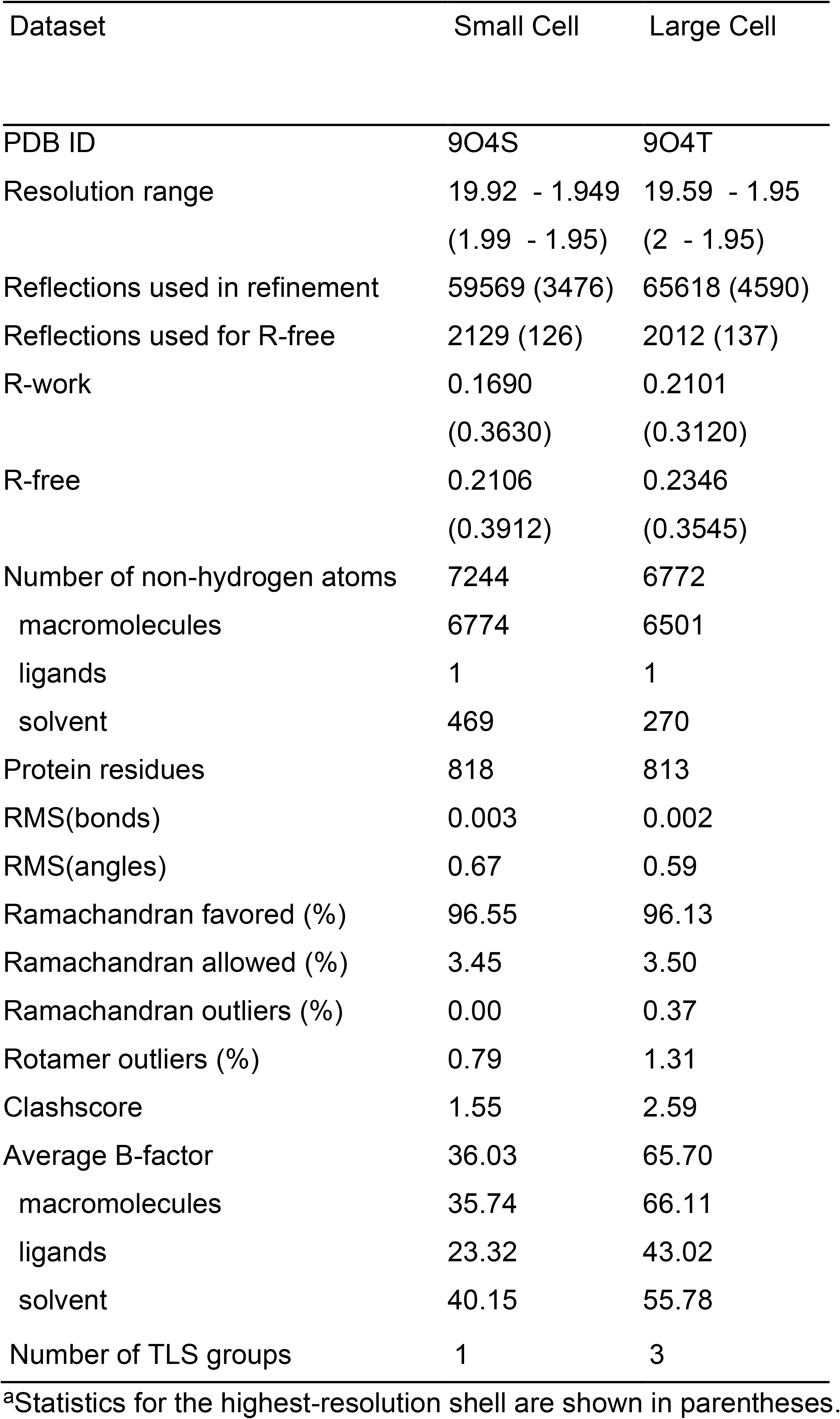
Model refinement statistics^a^.

### 2.7 Dehydration Experiments

Crystals were grown as described above, adjusting protein concentration from 15 mg/mL down to 4-6 mg/mL (0.2M NaOAc pH=5.5, 9% PEG-3350) to ensure growth of larger crystals. Individual crystals were harvested in a MiTeGen SLEEC humidity-controlled chamber and stored overnight in custom cassettes from MiTeGen. Crystals were mounted open to air under a humidity-controlled jet (Sanchez-Weatherby *et al*., 2009) at SSRL BL12-2. Relative humidity was set to 99.5%, a series of 10 images were collected using 0.5 ° oscillations for a total exposure of 50 kGy. The crystal was translated to avoid radiation damage, rotated to the original phi angle, humidity was adjusted to 98.5%, 5 minutes for equilibration was given, then 10 images were collected as above. This procedure was repeated for a descending range of relative humidities until the crystal no longer diffracted well. Data were indexed using *dials* and *Xia2*, and the corresponding unit-cell volume was calculated for each humidity setpoint.

## 3. Results

### 3.1 SFX Data Collection from SLO Microcrystals

We performed SFX on microcrystals of ferrous SLO at the MFX endstation of the LCLS. We prepared a slurry of SLO microcrystals using an established batch crystallization method (Wolff *et al*., 2020), starting from solution conditions that yield large, single crystals of SLO in vapor diffusion experiments (Minor *et al*., 1996). The batch-grown microcrystals had a rod-like morphology and were ∼20-50 μm in their longest dimension. These microcrystals were concentrated and mixed with a solution of 18% hydroxyethylcellulose dissolved in crystallization mother liquor, embedding them in a gel-like medium that was delivered to the X-ray beam using a high-viscosity extruder (Fig. 1a). SFX diffraction data were collected using the parameters described in Table 1.

**Figure 1.**
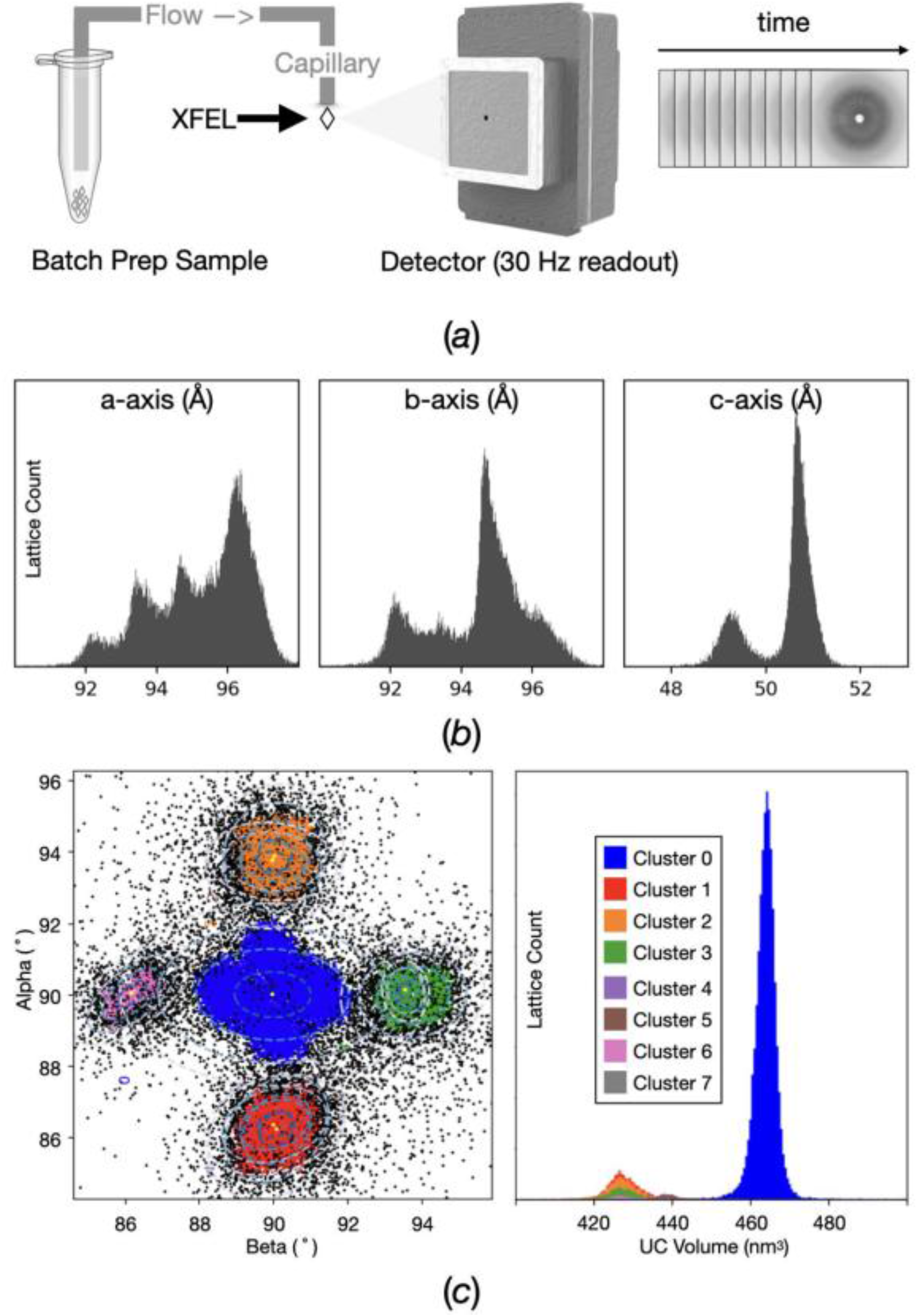
Unit-cell heterogeneity in serial crystallography data collected from SLO microcrystals. (a) Schematic of the serial crystallography experiment in which batch-prepared SLO microcrystals are delivered through a capillary to the XFEL interaction point until sufficient diffraction images are collected for a merged dataset. (b) Histograms of the refined unit-cell axes from approximately 66,000 indexed lattices reveal substantial heterogeneity in the crystal population. (c) DBSCAN clustering of the α, β, and c-axis unit-cell parameters identifies distinct lattice populations. Clusters are visualized by color in α–β space (left) and by unit-cell volume (right), demonstrating that the observed heterogeneity comprises discrete crystal forms with distinct unit-cell volumes rather than arising solely from indexing ambiguity.

### 3.2 Unit Cell Polymorphism Complicated by Indexing Ambiguities

During real time data analysis, after indexing approximately 66,000 lattices, we observed multi-modal distributions in each of the unit cell axis lengths (Fig. 1b). Because of the dynamic nature of serial crystallography experiments and the independence of each measurement, a variety of factors can affect the unit cell parameters measured for each crystal. Instrument factors such as photon energy variation or detector distance offsets are unlikely to account for the full magnitude of the observed differences, although small systematic contributions cannot be entirely excluded, so we began comparing unit cell distributions within and across sample batches (Fig. S1). The data were split into “runs” representing bins of sequentially-collected diffraction images, with each run representing approximately 5 minutes of data collection at 30 Hz repetition rate. Examining detector distance and the distribution of observed unit cell volumes as a function of data collection run revealed a bimodal distribution in unit cell volume that was not correlated to the refined detector distance (Fig. S1). The peaks of this distribution centered around two volumes, ∼460 nm^3^ and ∼420 nm^3^, which we will refer to hereafter as the “large-cell” and “small-cell”. Upon switching to a new sample batch, we saw a shift in the ratio of large-cell to small-cell lattices, but both were still present.

Next, we needed to partition the large-cell and small-cell diffraction patterns for downstream processing and merging. To achieve this, we turned to clustering methods, leveraging the DBSCAN algorithm (Ester *et al*., 1996). Upon exploring the six unit-cell parameters—*a*, *b*, *c*, α, β, and γ—it became clear that while the distribution of unit-cell volumes was bimodal, other factors were contributing to the observed unit cell heterogeneity (Fig. 1c). Specifically, the distribution of *c*-axis values was bimodal, matching the distribution of unit cell volumes, yet the *a*-axis and *b*-axis distributions were more complex, indicating potential indexing ambiguity. The potential for indexing ambiguity was further supported by consideration of the expected unit cell and space group. SLO crystal structures in the PDB (e.g. 5T5V) share a common crystal form.

This crystal form is monoclinic, with *P*2_1_ space group symmetry, however the *a* and *b* axes are approximately equal in length, and the β angle is nearly 90°, making the lattice pseudo-tetragonal. The consequence of this pseudosymmetry is that four different, but all valid, indexing solutions become difficult to distinguish based on unit cell parameters alone, and automated indexing algorithms struggle to apply a consistent indexing solution across all images in the dataset. In the case of SLO, if one chooses a “correct” indexing solution, then it is possible for the *a* and *b* axes to be chosen differently because they are similar in length but unique, for the direction of the c-axis to be given different polarity because the β angle is close to 90°, or for both of these mistakes to be made simultaneously. A similar indexing ambiguity in SFX has been reported previously (Hadjidemetriou *et al*., 2022), although in that case without additional unit-cell polymorphism.

To resolve the polymorphism and indexing ambiguity, we assigned all crystal lattices to space group *P*1, ignoring symmetry, and turned to a unit-cell clustering approach (Brewster *et al*., 2025). We used the α and β angles, allowing us to resolve a majority of the data into 8 clusters of interest, with a distinct 4-fold pattern. At first pass, we were able to separate and reindex 4 clusters (1, 2, 3, and 6 in Fig. 3C) corresponding to the small-cell, which were easier to separate due to larger deviations of the unit cell angles from 90°. We chose the cluster with α = 90° and β > 90° to be the reference, consistent with the expected monoclinic lattice, and applied one of three reindexing operators to each of the other clusters, implementing a consistent indexing solution across all images in the dataset corresponding to the small cell. On the second pass, we filtered data to isolate large-cell data (cluster 0 in Fig. 3C), then reclustered the lattices again based on α and β (not shown). With this second set of clusters, we resolved the 4 indexing solutions for the large-cell data. Consistent with the substantial difference in unit-cell volume between the two polymorphs, CCiso and Riso calculated between the large-cell and small-cell datasets (CCiso = 0.657, Riso = 0.541, overall, anisotropically scaled) confirm that the two datasets are strictly non-isomorphous. Our ability to sort the data into large-cell and small-cell polymorphs and solve a two-fold indexing ambiguity for each polymorph allowed us to determine two unique structures of SLO from a single sample of batch-grown microcrystals (Tables 2 & 3).

### 3.3 Unit-Cell Polymorphism is Correlated with Reduced Solvent Content

After determining structures for each of the two SLO polymorphs, we sought to understand the origin of the difference in unit cell volume. Given that protein crystals are composed of dynamic macromolecules and bulk water, both of which are subject to fluctuations, the shift in cell volume could have resulted from: 1) changes in the atomic packing of SLO or 2) changes in the intermolecular spacing of molecules in the crystal lattice. These factors are interdependent, so we aimed to determine the contribution of each to the overall loss of ∼8.5% of volume when moving from the large-cell to the small-cell (Fig. 2a). Meanwhile, the volume of a single SLO molecule derived from the large-cell and small-cell structures was effectively equivalent, suggesting there is no significant change in the overall density of atomic packing within individual SLO molecules. After accounting for both copies in the unit cell, the protein volume remains unchanged, and the remaining volume loss can be attributed to changes in the volume of bulk solvent within the crystal. This indicated that the primary driving force was a change in intermolecular spacing. Visualizing both copies of the ASU and the unit cell boundaries (Fig. 2b) further supported that the ∼4% decrease in the a-axis and ∼3% decrease in the c-axis (Table 4) were accounted for primarily by a decrease in intermolecular spacing. This observation prompted us to ask how the resulting crystal contacts were distributed and how this might impact SLO’s structure. We analyzed crystal packing interfaces for each of the two structures using PISA (Krissinel & Henrick, 2007). The interfaces with the greatest change in surface area were along the a-axis and c-axis (Table 5), which aligns well with those unit cell parameters being most affected (Table 4). Furthermore, the overall packing interface increased by over 58% in the small-cell as compared to the large-cell (Tables 4 & 5).

**Figure 2.**
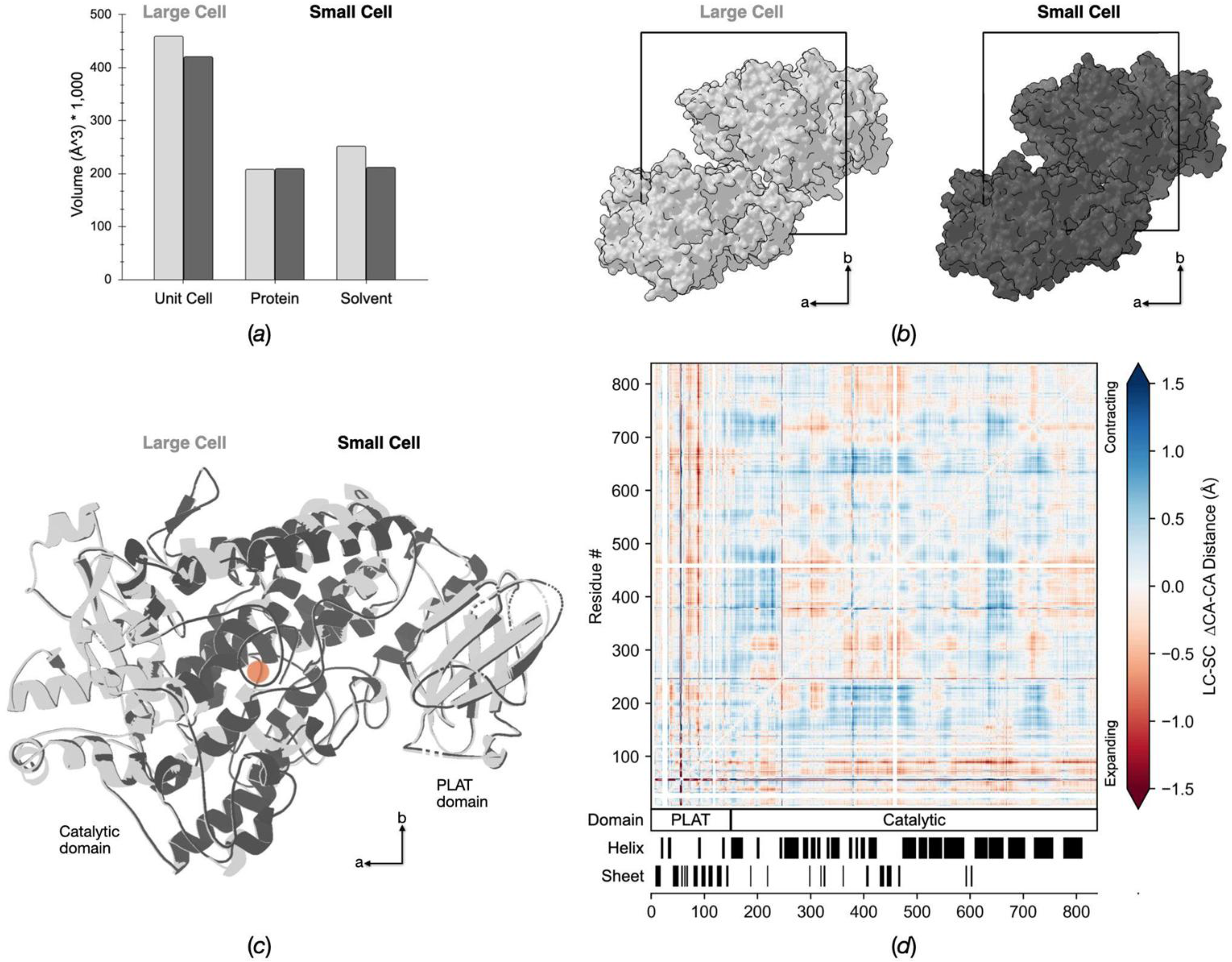
Origin of change in unit-cell volume and the resulting impact on SLO’s structure. (a) Quantification of volume of components filling the crystallographic unit-cell for both structures. (b) Visualization of the full unit cell for each structure, with protein copies shown as surfaces while the solvent is shown as whitespace, with the cell boundaries outlined in black. (c) Cartoon overlay of SLO as found in the ASU of both the large and small unit-cell structures, with the active site iron shown as an orange sphere. (d) A heatmap showing the internal CA-CA distance matrix of the small-cell structure subtracted from the internal CA-CA distance matrix of the large-cell structure reveals patterns of structural variation.

**Figure 3.**
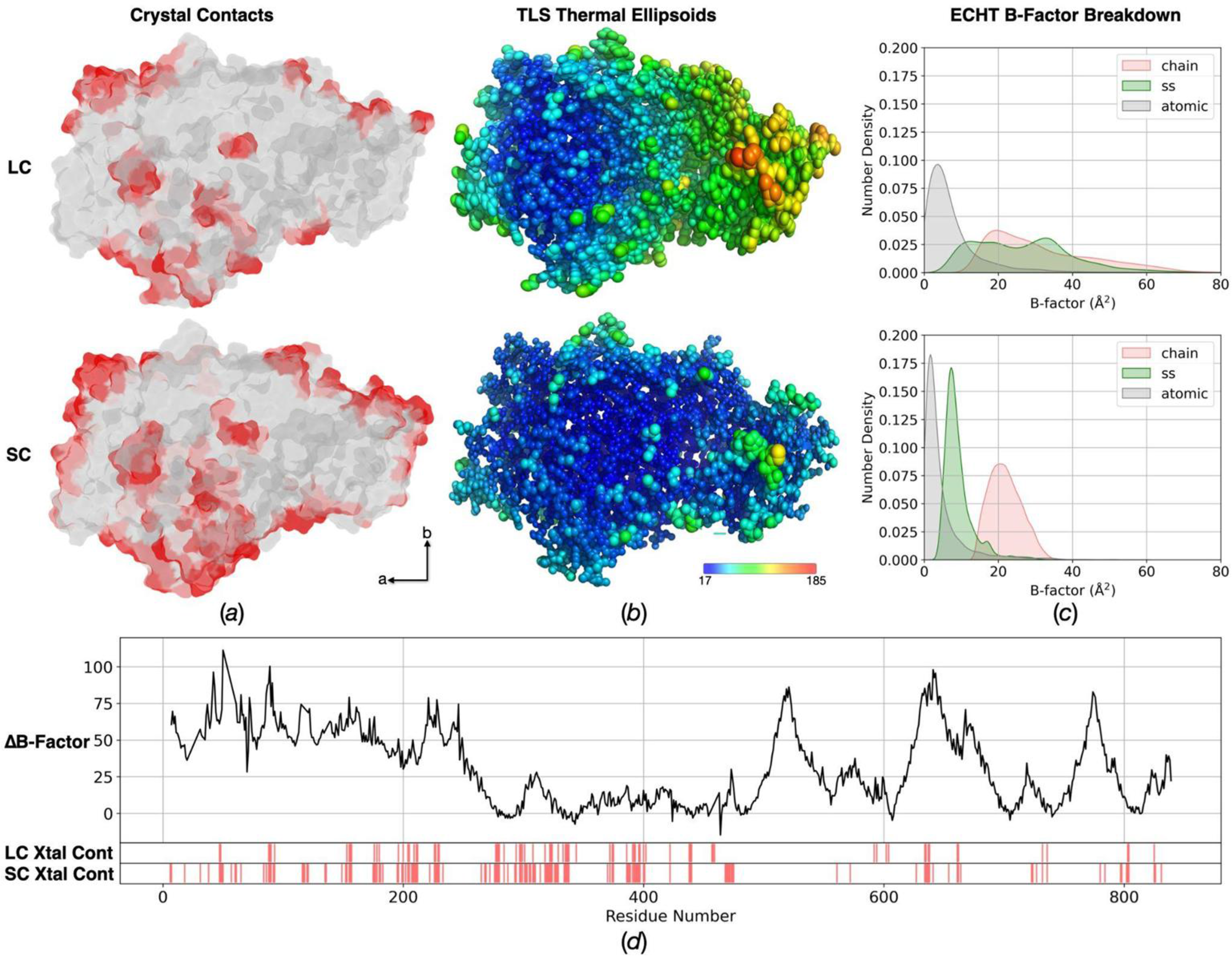
Impact of unit-cell shift on crystallographic B-factors. (a) Surface rendering of SLO with residues colored red that are designated as part of a crystal packing interface by PISA analysis. (b) TLS thermal ellipsoids are visualized for non-hydrogen protein atoms, with the ellipsoid’s size and color scaled to the ADP’s magnitude. (c) Density plot of the ADPs derived from PanDEMIC’s Extensible Component Hierarchical TLS (ECHT) analysis to breakdown contributions due to chain motions, secondary structure motions, and atomic fluctuations. (d) A line plot showing per-residues differences in total ADP values for LC-SC, with red bars below showing PISA-designated crystal contacts.

**Table 4.**
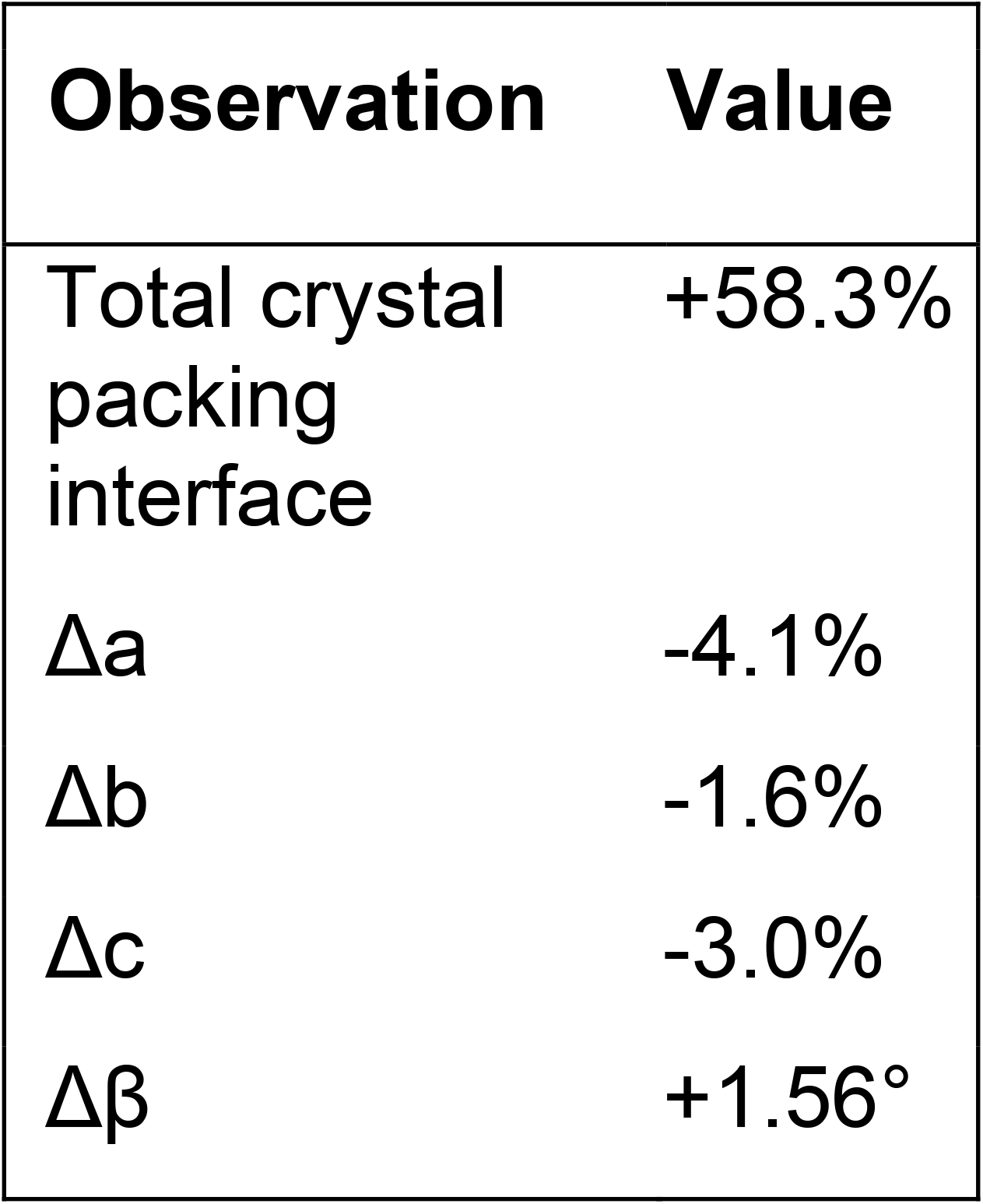
Summary of crystallographic changes from the large to small cell.

| Observation | Value |
| --- | --- |
| Total crystal packing interface | +58.3% |
| $\Delta a$ | -4.1% |
| $\Delta b$ | -1.6% |
| $\Delta c$ | -3.0% |
| $\Delta \beta$ | +1.56° |

**Table 5.** PISA analysis of crystal contact interfaces.

| Packing interface (symm operator) | Large cell (Å<sup>2</sup>) | Small cell (Å<sup>2</sup>) | $\Delta$ Interface (Å<sup>2</sup>) | UC parameter(s) most affected |
| --- | --- | --- | --- | --- |
| x-1,y,z | 23.5 | 400.8 | +377.3 | a-axis |
| x,y,z-1 | 248.6 | 574.8 | +326.2 | c-axis |
| -x+1,y-1/2,-z | 52.6 | 382.0 | +329.4 | a + c + $\beta$ shear |
| -x+1,y-1/2,-z-1 | 166.7 | 368.3 | +201.6 | a + c + $\beta$ shear |
| -x,y-1/2,-z-1 | 122.6 | 178.0 | +55.4 | preserved packing |
| -x,y-1/2,-z | 307.6 | 307.7 | +0.1 | negligible |
| **Total** | **921.6** | **2211.6** | **+1290.0** |  |

To demonstrate the coupling between solvent content and unit cell volume, we performed a crystal dehydration experiment. We used a large, single crystal of SLO and measured changes to the unit cell parameters as a function of relative humidity (RH) at a synchrotron beamline. We measured the unit cell volume as a function of RH and saw a trend toward smaller volume at lower RH (Fig. S2). Interestingly, in this humidity-dependent experiment performed on a large (approximately 50×50×200 μm^3^), single SLO crystal, the shift in volume did not reveal a sharp discontinuity as we expected based on the bimodal distribution of unit cells observed for cellulose-embedded SLO microcrystals (approximately 5×5×20 μm^3^), nor did this relationship follow a simple linear trend.

### 3.4 Coordinated Structural Rearrangements Result from Changes in Crystal Contacts

A global alignment of the large-cell and small-cell structures revealed an all-atom RMSD of 0.34 Å (Fig. 2c), indicating good agreement between the two models. However, global RMSD calculations are sensitive to alignment selection and can hide local changes, so we conducted a difference distance matrix (DDM) comparison to gain further insight into structural differences between polymorphs (Fig. 2d). Careful analysis of the DDM revealed that the majority of the PLAT domain (residues 1-145) moves away from the catalytic domain in the small-cell as compared to the large-cell structure. This is consistent with the greatest increase in a crystal interface being along the a-axis (Table 5), indicating that as contact is increased the PLAT is pushed down, away from the rest of the molecule. Immediately following the PLAT are short helical sections connected to several surface loops. This region (residues ∼151-240) and a nearby series of helices (residues ∼630-680) are pinned closer to the rest of the catalytic domain in the small-cell structure than in the large-cell structure. We further assessed this by mapping the residues designated as part of a crystal interface onto a rendering of SLO’s molecular surface (Fig. 3a). This revealed significantly more surface area involved in crystal contacts on the PLAT domain in the small-cell structure when compared to the large-cell structure. These coordinated, domain-level rearrangements, including repacking of the α-helical catalytic domain and a change in its positioning relative to the N-terminal PLAT domain, are not readily apparent from a simple structural overlay.

### 3.5 Differences in Crystal Packing Alter SLO’s Intramolecular Dynamics

Beyond changes in average coordinate position, we were curious how the crystal contacts affected the conformational ensemble of the protein. The observed shift in crystal packing from the large to the small unit cell was coupled to a decrease in overall crystallographic disorder, as evidenced by the decrease in Wilson B-factors from 32.41 to 23.15 Å^2^ (Table 2) as well as the decrease in the average atomic B-factors for protein atoms from 66.11 to 35.74 Å^2^ in the refined structures (Table 3). We refined our atomic models using a translation-libration-screw (TLS) model of anisotropic B-factors, and visualized the resulting thermal ellipsoids, which confirmed that rigid-body rotation of SLO within the crystal lattice, modeled using TLS, was significantly restricted in the small-cell (Fig. 3b). Because the TLS model imposes strict correlations in the ADPs of assigned rigid body groups, it can absorb (and thus incorrectly model) contributions to molecular disorder that occur on length scales that are smaller than the rigid group. To separate these contributions, we performed an algebraic decomposition of the anisotropic B-factor models derived from TLS refinement (Pearce & Gros, 2021). Disorder was modelled at three levels: 1) rigid body rotations and translations of the entire protein chain, 2) correlated movements of secondary structure elements, and 3) contributions from individual atoms. By maximizing the contribution at each level, then modelling the remaining disorder using the next level we determined that rotations of the entire chain were insufficient to explain the refined anisotropic B-factors (Fig. 3c and Fig. S3). Indeed, rigid body movements of the entire protein chain within the crystal lattice differed in magnitude and direction between the two hydration states, reflecting the observed shift in crystal contacts and packing around the PLAT domain. We observed that in the large unit cell, correlated movements of secondary structure elements (ss) account for a larger contribution to the anisotropic B-factor model, which we attribute to motions between the PLAT and catalytic domains. Visually, the directionality and magnitude of the thermal ellipsoids representing the anisotropic B-factors break across the domains, and there is evidence for more intramolecular disorder in the large-cell structure with higher solvent content. The decreased atomic B-factors of PLAT domain residues in the small unit cell correlates with increased crystal contacts across this domain (Fig. 3d). These trends in disorder are further corroborated by examining the location and distribution of ordered waters, and their average B-factors, which revealed increased order at the interface between the catalytic and PLAT domains in the dehydrated small-cell structure (Fig. S4).

### 3.6 Changes in Crystal Packing Distort the Interpretation of Side Chain Conformational Heterogeneity

Next, we were curious whether, in addition to the observed changes in atomic B-factors between the small and large unit cells, electron density maps corresponding to each of the two polymorphs would manifest differences in the appearance of discreet alternative conformations for amino acid side chains (Fig. S5). In the small cell, SLO has more surface residues displaying alternative conformations. Many of these residues are part of newly formed crystal contacts in the small-cell structure, highlighting how crystal packing can stabilize alternative side chain conformations at an interface, but residues buried in the core of the protein also display differences in the appearance of discreet side chain conformations (Fig. 4). Notable examples of such discrepancies include residues LEU546 and ILE552, both buried hydrophobic residues. In the large-cell structure, only a single rotamer is evident in the electron density, but the density is more diffuse (Figs. 4a & 4d) and the refined B-factor for the side chain atoms is higher. In the small-cell structure, the electron density contains evidence for two rotameric states that can be modeled discreetly, and their occupancies refined (Figs. 4b & 4e). Consistent with this observation, modeling a single rotamer for each of these sidechains results in residual mF_o_ -DF_c_ difference electron density for the small cell, but not for the large cell (Fig. 4a-d). Ringer analysis (Lang *et al*., 2014) provides additional support, revealing the clear presence of multiple peaks in the 2mF_o_ -DF_c_ electron density maps corresponding to alternative rotamers for these residues in the small-cell but not in the large-cell (Figs. 4c & 4f). The impact on the interpretation of conformational heterogeneity is notable, given that the same resolution cutoff was applied to each data set and the data quality indicators are similar for the two data sets (Table 2).

**Figure 4.**
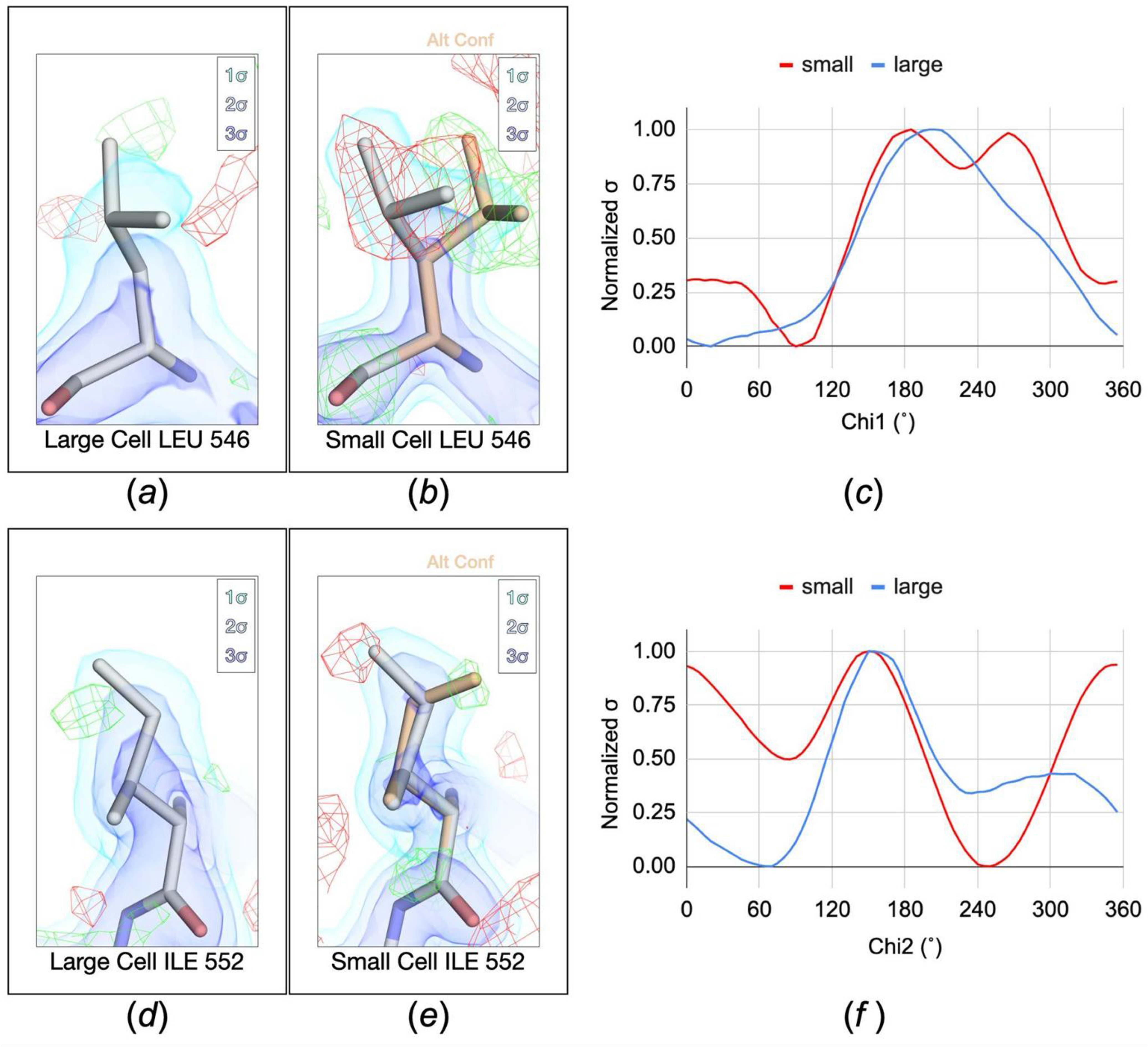
Electron density and modelling of alternative conformations differ between the large and small cell structures. (a, b, c, d) 2mF_o_ -DF_c_ electron density maps are carved within 2.0 Å of atoms, rendered as volumes contoured at 1σ (light blue), 2σ (blue) and 3σ (dark blue). (a,d) Large-cell structures modeled in a single conformation (grey sticks) with standard mF_o_ -DF_c_ difference density shown as green (+2σ) and red (-2σ) mesh. (b,e) Small-cell structures depicting major (grey sticks) and alternate (wheat sticks) conformations; mF_o_ -DF_c_ difference density (mesh) corresponds to an omit map calculated excluding the alternate conformation. (c, f) Ringer 2mF_o_ -DF_c_ density profiles sampled around the side-chain dihedral angles for Leu546 chi_1_ (c) and Ile552 chi_2_ (f), comparing the large-cell (blue) and small-cell (red) structures.

## 4. Discussion

In this work, we examined SLO microcrystals using SFX and found that our samples contained a mixture of two polymorphs, allowing us to determine two unique crystal structures with the same space group but different unit cell volumes. Analysis of the diffraction data was complicated by the presence of pseudosymmetry, caused by the *a* and *b* axes of the monoclinic unit cell being nearly identical in length, and the *β* angle being almost exactly 90°. We sorted the crystal polymorphs based on their unit cell volumes and applied a uniform indexing solution to the individual lattices based on a clustering approach (Brewster *et al*., 2025). The main difference between polymorphs is a difference in solvent content. While the all-atom RMSD was small (0.34 Å), careful analysis of structural differences revealed coordinated changes in position between the PLAT and catalytic domains. Additionally, TLS refinement showed that the change in crystal packing between polymorphs affects the interdomain flexibility of SLO in the crystal. Furthermore, the interpretation of side chain rotamer placement and occupancy in the core of the protein based on the electron density maps differs between polymorphs. This study highlights an underappreciated interplay between experimental and analytical challenges in macromolecular crystallography.

Our data illustrate two consistent complications that are inherent to serial measurements: crystal pseudosymmetry and sample polymorphism. Issues arising from pseudosymmetry, where the crystal lattice has higher symmetry than the true space group, were appreciated early on in the development of serial crystallography, and led to the development of clustering algorithms to unravel the “computational twinning” that would arise from applying an inconsistent indexing solution to individual lattices within the data set (Brehm & Diederichs, 2014; Gildea & Winter, 2018). Furthermore, other serial crystallography studies have also reported unit cell polymorphism that is similar to (and in some cases, even more complex than) what we observe in our data (Ebrahim *et al*., 2019; Stagno *et al*., 2017; Brewster *et al*., 2025; Hadjidemetriou *et al*., 2022). In our case, the presence of two very clear peaks in a histogram of unit cell volumes revealed the existence of two specific polymorphs, and allowed efficient sorting of the lattices into two groups. Observing the symmetry of *α* and *β* angle distributions in our data (Fig. 1c) suggested that there could be an indexing ambiguity, and consideration of the expected unit cell axis lengths and angles confirmed the possibility of pseudosymmetry and allowed us to determine a suitable set of reindexing operators. After developing an appropriate workflow for data processing that included sorting and reindexing, we were able to determine structures for each polymorph.

We next considered the nature of the polymorphism itself, with the main difference being that the smaller, more compact unit cell has less solvent and more extensive crystal contacts between neighboring SLO molecules. Discontinuous, humidity-dependent phase transitions involving changes in crystal solvent content have been reported for other macromolecular crystals, including hemoglobin (Huxley & Kendrew, 1953) and lysozyme (Dobrianov *et al*., 2001), and provide possible context for interpreting our observations, though we do not have sufficient data to establish that the polymorphism reported here reflects the same underlying phenomenon. While our single-crystal dehydration experiments establish a correlation between crystal hydration and unit cell dimensions, it is unclear why these measurements did not reveal a sharp discontinuity in unit-cell dimensions as a function of relative humidity (Fig. S3), mirroring the bimodal distribution of unit cell parameters observed in the serial experiments. Dobrianov, et al. noted that crystal size and shape can play a role in equilibration times for phase transitions in macromolecular crystals, and that these transitions can occur inhomogeneously (Dobrianov *et al*., 2001), which could explain why we observe complex behavior from a large single crystal, vs. a distinct shift in the distribution of unit-cell parameters for microcrystals. Previous work in serial crystallography also supports the notion that small crystals are more robust to changes in unit cell dimensions induced by experimental perturbations (Stagno *et al*., 2017).

An analysis of anisotropic B-factors refined using TLS parameters clarifies how changes in crystal packing and solvent content affect the conformational flexibility of SLO within the crystal lattice. We observed increased B-factors in the PLAT domain of the large cell model, which cannot be explained by rigid-body displacements of the crystallized molecules alone. Instead, anisotropic B-factors from TLS refinement suggest that the crystal packing in the large cell is more permissive of collective motions that alter the relative positioning of the catalytic and PLAT domains, comparable to the rocking motion suggested by solution scattering measurements on SLO (Dainese *et al*., 2005) and homologous lipoxygenases (Hammel *et al*., 2004). This is consistent with the observation that more extensive crystal contacts present in the small unit cell push on the PLAT domain, moving it away from the catalytic domain and restricting its mobility. Currently, it remains unclear which polymorph best represents the physiologically relevant ensemble of SLO, and the functional relevance of putative motions resolved in both the large and small cell structures is also unknown. Nevertheless, these observations illustrate how crystal packing and solvent content can alter the conformational flexibility of crystallized proteins.

In addition to differences in interdomain dynamics, we also noted an interesting difference in the manifestation of alternative side chain conformations in the core of the protein, exemplified by residues L546 and I552, which are both of functional significance. Notably, these residues have been implicated, via mutagenesis, in long-range motions that are coupled to catalytic turnover (Offenbacher *et al*., 2017b; Zaragoza *et al*., 2023). In the large cell, electron density maps reveal only a single side chain conformation for these two residues, but their refined B-factors are substantially elevated relative to the small cell. In the small cell, two distinct conformations can be modeled in the electron density, each with lower B-factors than the single conformations modeled in the large cell. These apparent differences in the electron density exist, even though the two structures were refined against diffraction data with a similar resolution cutoff. This observation raises the question of whether the crystal packing changes the nature of side chain motions or alters our ability to faithfully observe them. Specifically, does the larger unit cell promote harmonic movements of these atoms while the small cell promotes anharmonic exchange between rotameric wells? Or does the overall increased heterogeneity inherent to the large cell (Wilson B-factor of 23.15 Å^2^ versus 32.41 Å^2^ for the small cell) lead to a blurring of the electron density that simply eliminates our ability to resolve the alternative conformations of these sidechains? These questions are relevant, as the observed effects occur in a region of established functional importance. Along the same lines, the small cell model has significantly more ordered water molecules than the large cell model, despite being of a similar resolution. We likewise cannot determine whether this is because the enzyme interacts with fewer water molecules in the more expanded crystal lattice, or whether our ability to resolve them is compromised by increased overall disorder.

Although we could demonstrate the effect of crystal polymorphism on SLO’s dynamics, the cause of this polymorphism remains unclear. Based on the present data (Fig. S1), we cannot distinguish between the possibility that both polymorphs were already present in the crystal slurry prior to embedding, or the possibility that the crystals undergo a phase transition upon exposure to cellulose or pressurization in a high-viscosity microfluidic extruder. It is noteworthy, however, that other reports, including some of our own work, show that use of viscogens correlated with smaller dimensions of the crystal lattice. For example, CypA microcrystals measured in viscous hydroxyethyl cellulose show a 2% reduction in unit cell volume relative to the same crystals measured in a sample delivery medium lacking the viscogen (Wolff *et al*., 2020). Additionally, SFX experiments on photosystem II revealed that increased PEG concentrations lead to a 6% decrease in unit cell volume, which resulted in significantly lower Wilson B-factors and increased resolution (Young *et al*., 2016). These precedents are consistent with, though do not on their own establish, a role for the cellulose carrier medium in the polymorphism we observe in SLO.

Crystallographic experiments (including time-resolved SFX) are increasingly employed to explore the dynamics of crystallized proteins, and our study underscores the importance of considering how protein motion can be impacted by the crystal environment. At first, the difference between the two polymorphs present in our microcrystal samples seemed inconsequential - despite a difference in solvent content, the crystal resolution and data quality were comparable, and the refined atomic coordinates appeared similar. A more thorough inspection revealed important differences in the enzyme’s internal dynamics between crystal forms. Specifically, algebraic decomposition of the resulting anisotropic B-factor model allowed us to dissect rigid-body (i.e. crystalline) disorder from disorder resulting from intramolecular dynamics, which would not be possible from a simple, direct comparison of refined B-factors. This analysis shows that crystal solvent content is coupled to intramolecular flexibility, an observation that could have important implications for the understanding of SLO’s functional motions.

## Acknowledgments

The authors would like to thank Alexander Batyuk, Mark Hunter, and the MFX staff for support during experiments at LCLS.

## Author Declarations

### Author Contributions

**AMW, SC, JPK, ARO, FP, MAW, and MCT** conceptualized the experiments. **AMW, DWP, IDY, RGS, JPTZ, SC, AHF, ARO, NKS, MAW, and ASB** contributed resources and methodology. **AMW, DWP, IDY, AD, VG, JH, AH, RL, SL, RLM, DM, STMM, AO, JMR, SR, DMT, AHF, FP, ASB, and MCT** conducted investigations. **AMW, DWP, IDY, VG, DMT, AHF, ASB, and MCT** analyzed data. **JPK, FP, NKS, ASB, and MCT** provided project administration and secured funding. **AMW, AD, AH, ARO, and MCT** wrote the manuscript. **AMW, IDY, AD, AH, STMM, ARO, NKS, MAW, and MCT** edited the manuscript.

### Data Availability

Structural models and associated reflection data are deposited in the PDB under accession codes 9O4S & 9O4T. The model used for molecular replacement is available under code 4WFO. Raw diffraction frames are deposited in CXIDB under 242 (Wolff & Thompson, 2026).

