## Supporting Information: Figures for "Conformational flexibility of soybean lipoxygenase is coupled to crystal solvent content in serial crystallography"

Alexander M. Wolff, et al.

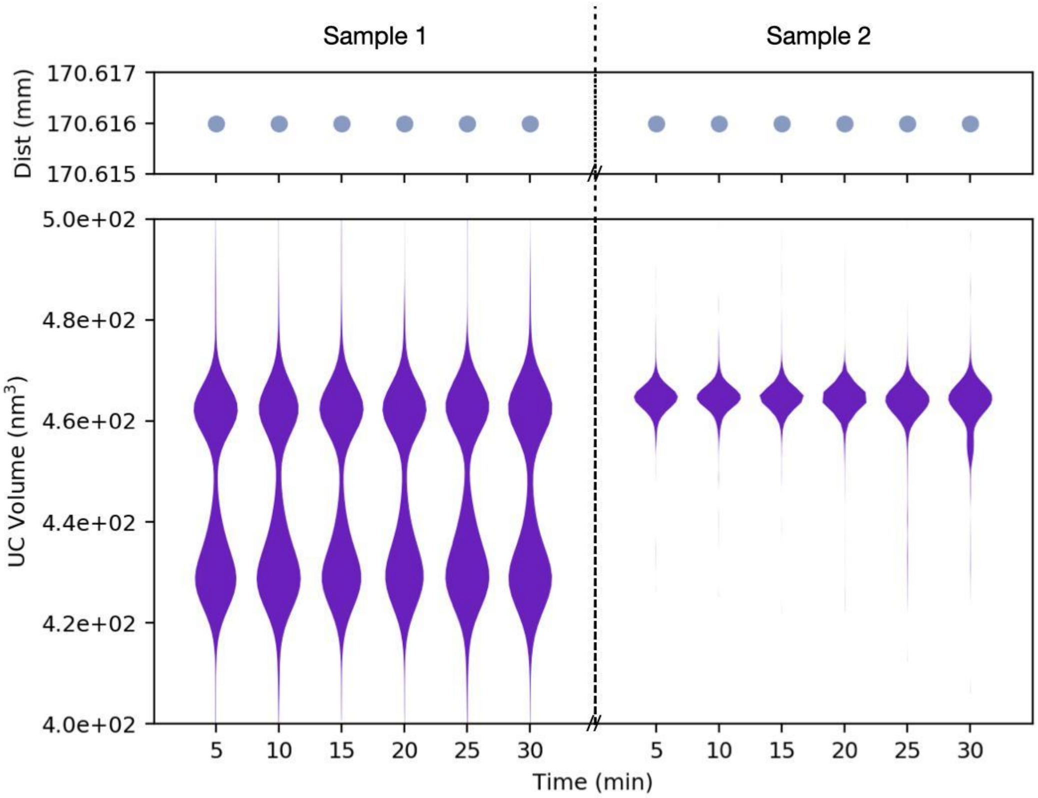


**Figure S1.** Unit cell heterogeneity in serial crystallography data collected on SLO. Unit cell volume violin plots (bottom) and detector distance scatterplot (top), displayed on a per-run basis where each run corresponds to ~9,000 images, collected over 5 minutes, with a sample change denoted by the dotted vertical line and corresponding break in the horizontal axis.

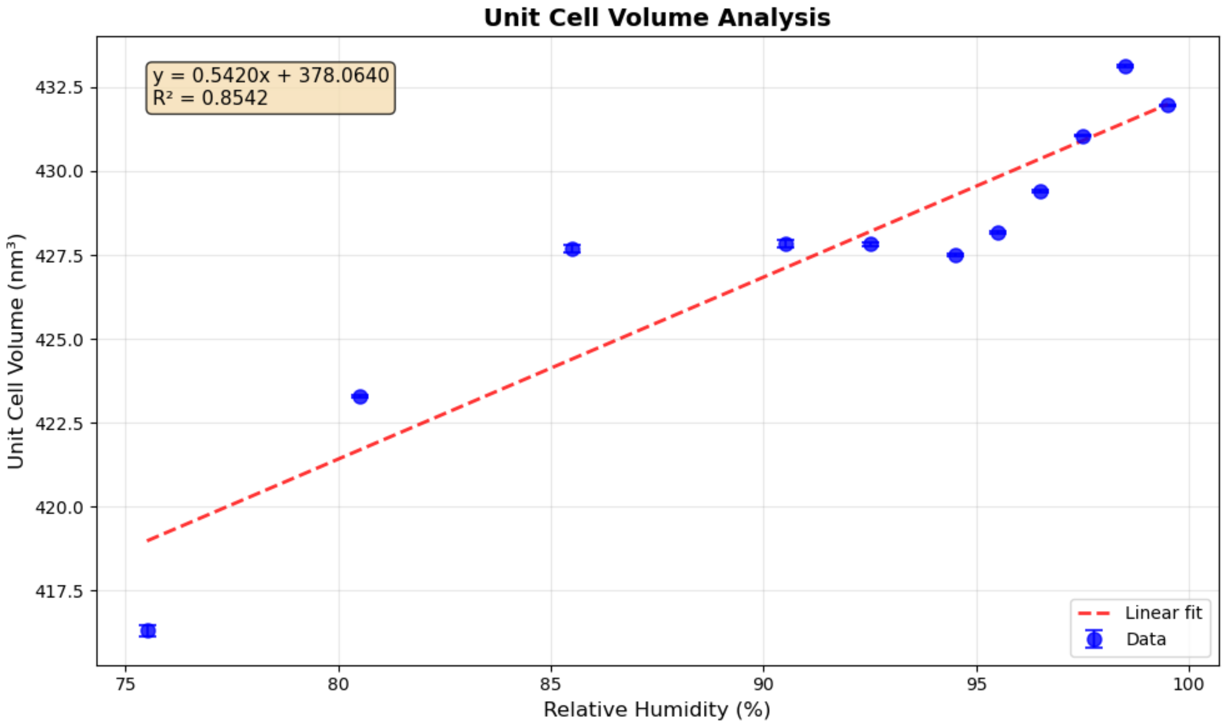


**Figure S2.** Hydration dependent changes in unit-cell volume of a single SLO crystal. Calculated unit-cell volume is shown based on indexing results from a series of 5-degree wedges collected from a single SLO crystal. Hydration was controlled using a humidity jet, beginning at 99.5% RH, following a change in the RH setting the crystal was given 5 minutes to equilibrate before collecting the subsequent wedge of images to assess the unit-cell volume.

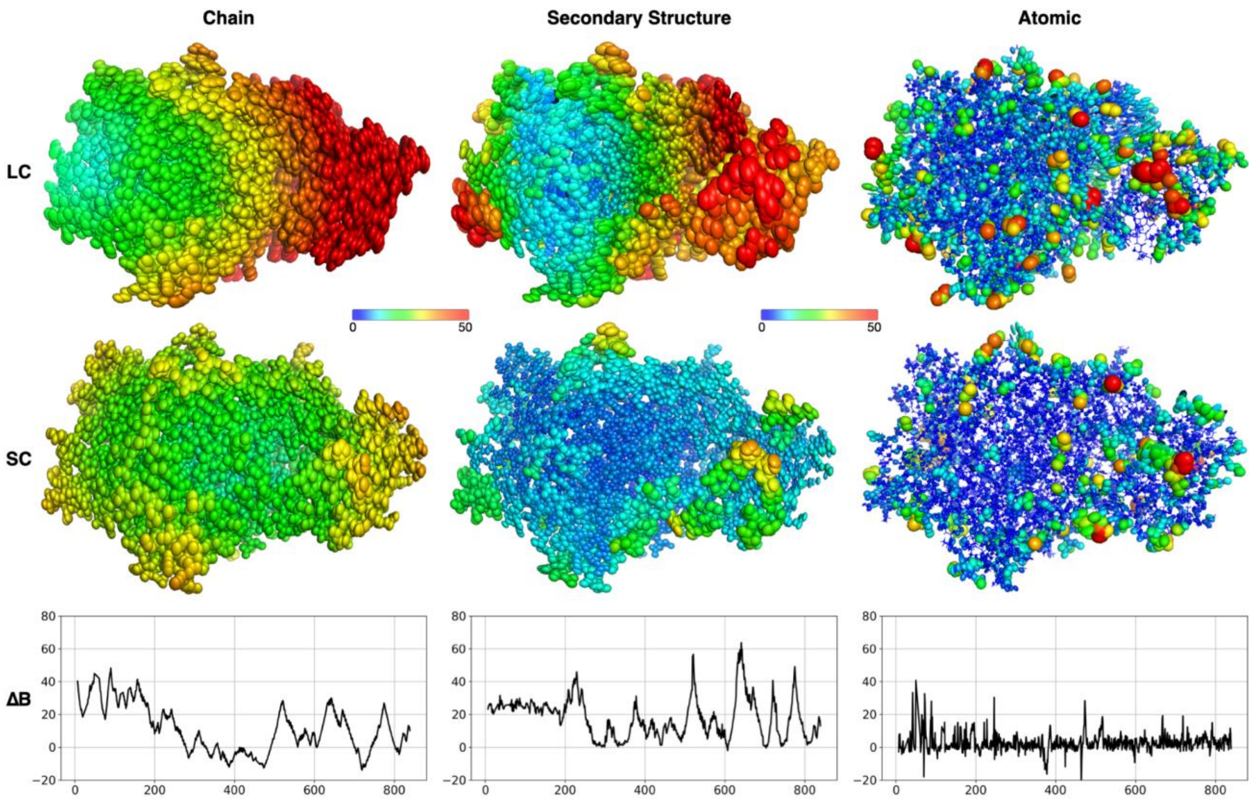


**Figure S3.** ECHT B-Factor Differences Between Hydration States. TLS thermal ellipsoids are visualized for non-hydrogen protein atoms, with the ellipsoid's size and color scaled to the ADP's magnitude. ADPs from the full model were separated using PanDEMIC to break down contributions due to chain motions, secondary structure motions, and atomic fluctuations, with each component visualized independently. In the bottom row the difference between the B factors (large cell - small cell) is plotted on a per-residue basis.

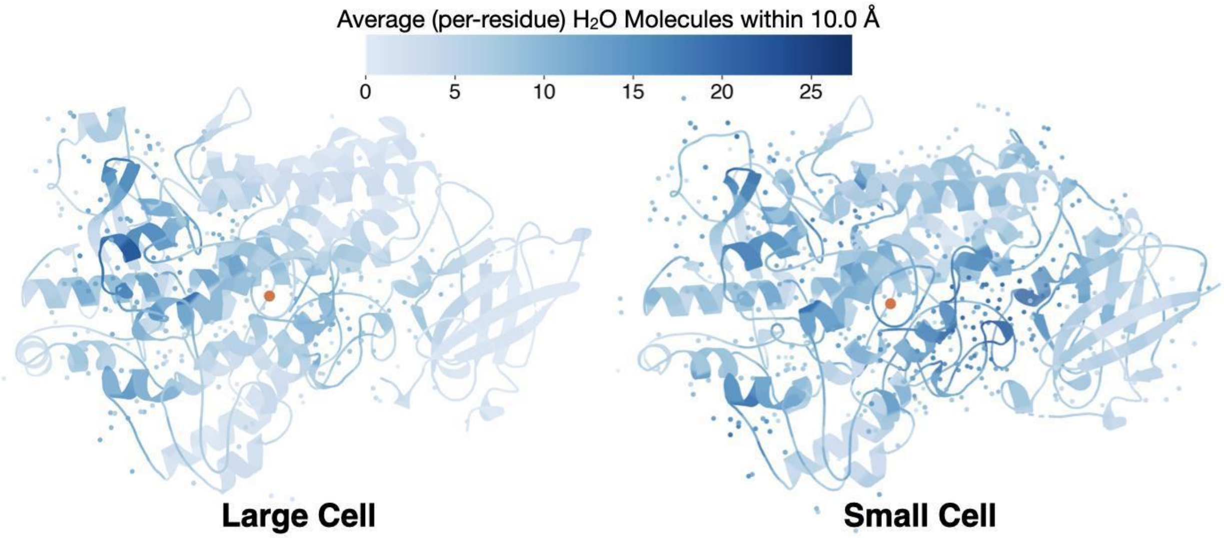


**Figure S4.** Distribution of Ordered Waters. SLO's structure is illustrated as a cartoon, with the active site iron shown as an orange sphere and ordered waters shown as smaller spheres. The cartoon and ordered waters are colored based on the average count of $H_20$ molecules within a 10 Å radius of the residue, highlighting that there are more ordered waters in the small cell structure, which are also distributed differently across the molecule.

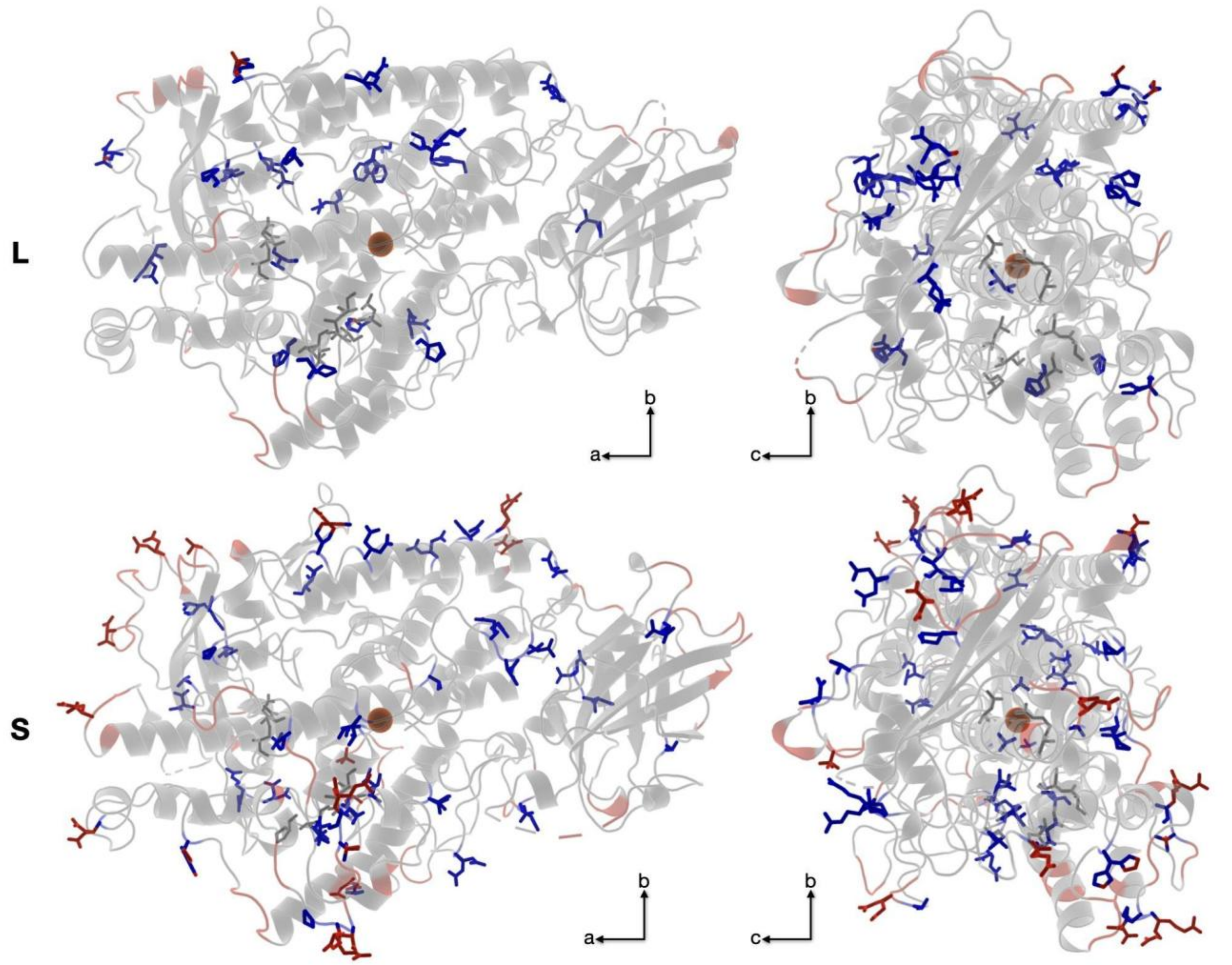


**Figure S5.** Distribution of Alternative Conformations. SLO's structure is illustrated as a cartoon, for the large cell (top panels) and small cell (bottom panels), with the active site iron shown as an orange sphere and residues with alternative conformations modeled shown as blue or red sticks. Blue denotes alternative conformations only, while red denotes crystal contacts. There are a greater number of alt confs in the small cell structure, with regional disparities focused near the active site.
